# Effects of Transport and Altitude on Hormones and Oxidative Stress Parameters in Sheep

**DOI:** 10.1101/2020.12.21.423758

**Authors:** Hilal Tozlu Çelık, Fatih Ahmet Aslan, Diler Us Altay, Metehan Eser Kahvecı, Kalbiye Konanç, Tevfik Noyan, Sertaç Ayhan

**Author notes:** Corresponding author, –.

## Abstract

With this study, it was aimed to determine the stress effects that can be caused by transporting and altitude in sheep. Karayaka sheep were used in the study. The live weight of the sheep (n=30) while hungry was determined before transport and sea level. Average live weight was determined as 55.64 ± 4.66 kg. Blood samples were collected just before (sea level) and just after transport (1500 meters above sea level). Transportation distance was approximately 182 km and duration was 5 hours. According to the findings, cortisol was not affected by transport stress and altitude (p>0.05) and Triiyodotironin (T3) (p<0.039) and Tyrosine (T4) (p<0.000) were affected significantly. Malondialchehyche (MDA), which is one of the oxidative stress parameters, was significantly affected (p<0.039) and Protein Carbonyl (PC) values were not affected by transport and altitude (p>0.184). As a result of this study, it was determined that transportation and altitude in sheep causes stress. Stress-reducing measures should be taken in the exposure of sheep to altitude differences and in transportation. Antioxidant nutritional supplements should be made in order not to negatively affect the meat quality in sheep.

## Introduction

There are many effects that cause stress in sheep breeding. These are veterinary procedures (vaccination, surgical intervention, treatment and blood tests), animal breeding practices (weaning, shearing, nail cutting, numbering), adverse weather conditions (extreme heat and cold) and malnutrition [1]. In addition, during the transportation of animals, bad road conditions, driving performance of the drivers, transportation vibration, road length, climate, lack of water and food can cause stress in animals. Stress can cause biochemical and physiological effects in animals [2–3–4–5–6]. In the metabolic processes in the cells of aerobic organisms, toxic free radicals appear during the reduction of oxygen to water. Free radical sources in living systems can emerge during the organism’s own metabolism, or they can enter the living organism in various ways. Free radical types can become stable by taking electrons from molecules such as lipids, proteins, carbohydrates, nucleic acids [7]. Although free oxygen radicals play a role in very important processes such as mitochondrial respiration, platelet activation, leukocyte phagocytosis and prostaglandin synthesis, they have toxic effects on lipids, proteins, DNA and carbohydrates. Therefore, it can cause structural and functional disorders in cells [8–9]. Free radicals cause lipid peroxidation in lipids [10], resulting in membrane damage. One of the products formed because of free radicals peroxidation two or more unsaturated fatty acids in lipids in cell membranes is MDA. This molecule is an indicator of oxidative degradation and tissue damage of lipids. Because of protein oxidation is formed in end products such as protein carbonyl, oxidation of nitro tyrosine and thiol groups. These products are indicators of protein oxidation [11–12]. They react physiologically to the changing environmental conditions. These reactions occur in the form of differences in body temperature, T3, T4 hormone levels, respiratory and pulse rates [13]. It has been reported that there is a great increase in cortisol and prolactin concentrations at the beginning of the journey during transportation in sheep and changes in hormone release within the first 3 hours of the 15-hour journey [14]. Minimizing the stress that may occur during the transportation of animals is important for animal health and meat quality. Therefore, reducing stress in animals will prevent economic losses for the breeder [3–15–16–17]. The productivity of sheep is affected by hypoxia and oxidative stress at high altitudes due to altitude difference. The development and functions of the corpus luteum can be affected and the reproductive performance of sheep may be reduced at high altitudes [18]. It was determined that the red blood cells of sheep and goats raised at high altitude (over 1000 meters) were higher than those raised in low altitude regions, and it was determined that age, gender and altitude affect the morphology of red blood cells [19].

Sheep breeding in the coastal region of the Black Sea region is carried out in certain periods of the year as a migration to the plateau. Migration to plateau takes place twice a year in the region. Altitude and distance are quite high between at the sea level and the plateau. Two methods are used to transportation the sheep to the plateau. First method, the sheep is transported by walking. The second method is transportation by transport. In recent years, mostly trucks have been used for transporting sheep to plateau. Environmental impacts such as transportation conditions of the sheep, engine noise, movement and stopping of the sheep during the transportation with the truck can create stress in the sheep. With this research, the physiological effects and oxidative stress effects that may occur in sheep transport and altitude were tried to be determined. This study is important to give an idea to minimize the negative effects that may occur with transport and altitude in sheep.

## Methods

### Animal material

The sheep pen where the research will be applied, was decided. It is aimed to determine the stress effects that may occur during by transport to the Perşembe plateau (1500 m) of Karayaka sheep. They are transported from Tekkeköy (sea level) to the Perşembe plateau as of the end of April and the first weeks of May in the Black Sea region. For this reason, the necessary materials for the research were taken and the study started on May. The livestock farm in Tekkeköy where the sheep are located, has been visited. It is a settlement in Samsun province (Samsun-air temperature 27 ° C-Humidity 70%). The sheep were transported to the Persembe plateau (17°C-Humidity 28%) at 1500 altitude above sea level. It is located in Aybastı district of Ordu province. Both places are located in the north of Turkey and the Black Sea climate is dominant (winters are cool and humid and summers are warm and humid). The transport distance is approximately 182 km. A truck was used to transport the sheep. A total of 30 healthy Karayaka sheep (approximately 2 years old) were used. The sheep of same age were selected and their ear numbers were recorded. They were marked with harmless paint. In the research, only female animals were used. Weighing was carried out early in the morning while the animals were hungry and before transportation (sea level). Feed and water were given before transport in the sheep. The feeding arrangement applied by the breeders has not been changed. The sheep are placed in the compartments separated in the transport in order to prevent accumulation in the transport. The transport stopped every 2 hours and the condition of the sheep was checked. They were transported by truck for 5 hours. Average speed of the truck was 46 km/h. Feed and water were not given during transportation.

### Blood sampling, Biochemical parameters and hormone monitoring

Blood samples were collected from each weighed animal for biochemical parameter and hormone analysis. These treatments were carried out two times, before the sheep were put into the transport (sea level) and after the transportation (1500 m) was completed. The sheep were rested after transport (1500 m). Blood samples were taken again. 10 ml of blood samples were taken from each sheep’s jugular the vein [13]. The blood samples taken into gel biochemistry tubes were centrifuged for 10 minutes at 3000 r.p.m after approximately 30 minutes of waiting time and their serums were separated. The collected serum samples were dispensed into 1.5 ml Eppendorf tubes. The collected blood samples were quickly kept in ice pack with car refrigerator and sent to the laboratory and stored at −24 °C. Separated serums were sent to Ankara Düzen Laboratory for T3, T4 and Cortisol hormone analysis and analyzes were performed (ECLIA (Roche)) and the results were obtained. The blood samples taken were sent to Ankara Düzen laboratory in cold chain.

MDA and PC values of oxidative stress parameters in sheep were analyzed by using Sheep Malondialchehyche Elisa Kit (English Instruction-enzyme-linked immunosorbent assay-Catalogue No:201-07-0086) and Sheep Protein Carbonyl Elisa Kit (English Instruction-enzyme-linked immunosorbent assay-Catalogue No: 201-07-0098). MDA and PC were analyzed in Ordu University Medical Faculty Biochemistry Laboratory and their results were obtained.

## Data analysis

The normal distribution control of the data was checked with the Kolmogorov-Smirnov test. Homogeneity control of group variances was done by Levene test. Introductory statistical values of variables (property=variable) such as mean, standard error, standard deviation, coefficient of variation, minimum and maximum values were calculated. Paired t test was applied and Pearson correlation test was performed to determine the relationships between the variables. The significance level (α) was taken as 5% in calculations and interpretations.

## Ethics statement

This research was found in accordance with the ethical principles and rules according to the decision number 10 taken by Ordu University Animal Experiments Local Ethics Committee at its meeting number 03 dated 19/07/2018.

## Results

### Body weight

The study was carried out in line with the scope and purpose specified in the project. The method was applied as decided in the project proposal and data were obtained. Sheep were weighed while hungry before the transport (sea level). As a result of the statistical analysis of the data, the average overall body weight for sheep (n=30) was 55.64 ± 4.66 kg (Table 1).

**Table 1.**
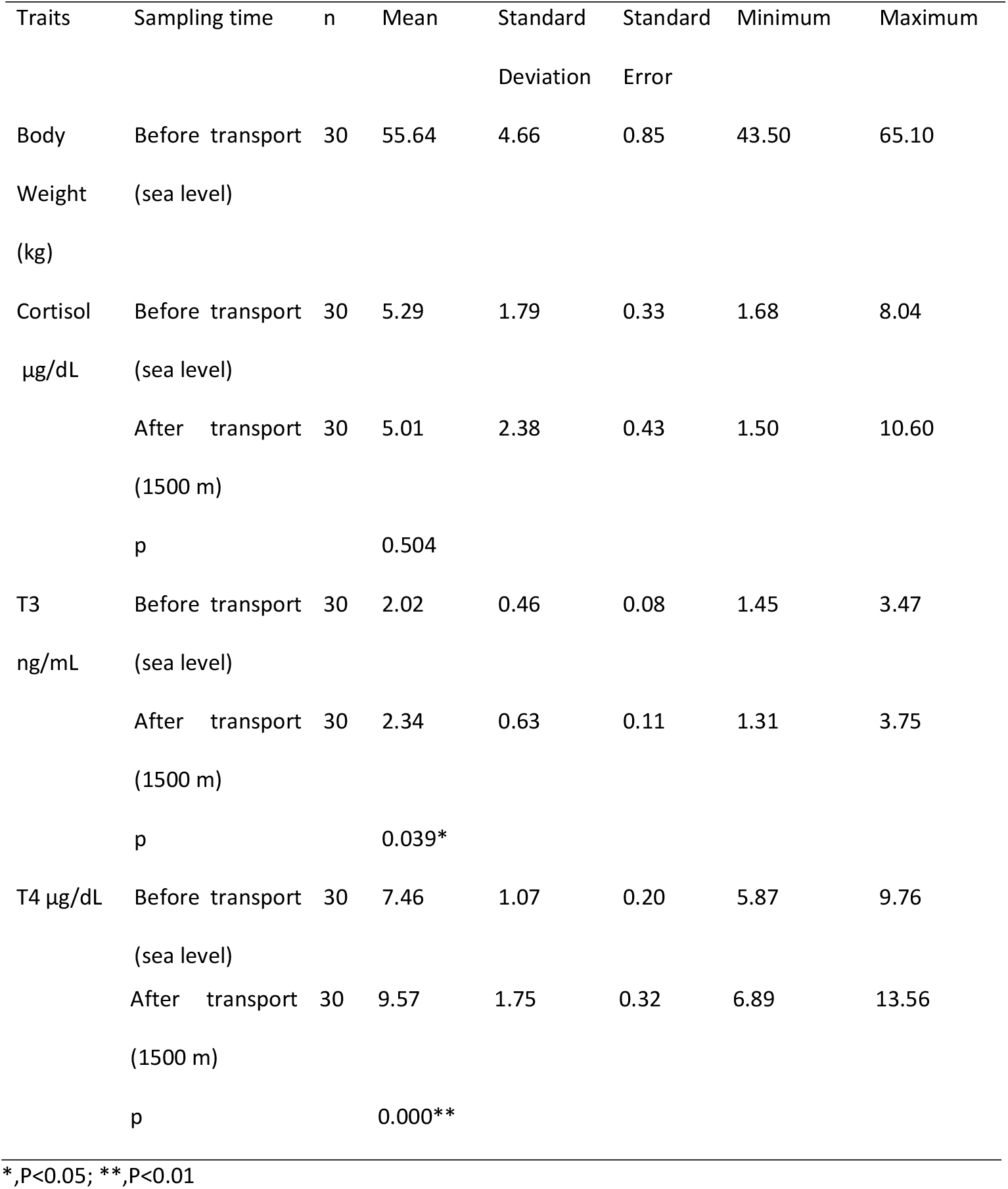
Hormone levels at different times in sheep (Cortisol, T3 and T4)

### Biochemical parameters and hormones

It was found that cortisol hormone was not affected by transport and altitude in the sheep (p=0.504). However, it is seen that the cortisol values are similar at the time of measurement. T3 hormone was significantly affected (p<0.039) and T4 hormone was highly significantly affected (p<0.001) (Table 1). After the Karayaka sheep were brought to the plateau (1500 m), the hormone T3 and T4 levels increased. The air temperature of the plateau is lower than the sea level. The findings are consistent with the report that the hormones T3 and T4 increase with decreasing temperature and altitude. The expected increase in cortisol level was not occur, but an increase in T3 and T4 hormone levels was occurred. MDA, which is an indicator of lipid peroxidation, was significantly affected by oxidative stress parameters (p<0.039) (Table 2). While the MDA level is expected to increase as a result of oxidative stress after transport (1500 m), it was found to be 1.80 nmol/L lower than the value before the transport. There were no statistically significant changes in protein carbonyl levels (p=0.184). However, while it is expected to increase with by transport, it was found 8.34 ng/ml lower than the PC level before the transport. According to these results, the fact that the level of MDA and PC before the transport (sea level) was higher than after transport (1500 m) shows that the sheep were also exposed to different stress factors before transport (sea level). Before transport, It was an extremely hot environment even though it was May in Black Sea Region. It can be said that Karayaka sheep are more sensitive to heat stress. It was observed that stress increased in sheep with the effect of humidity and temperature especially at sea level. In this direction, recommendations were made to improve the pen conditions of the breeder. A very significant (p<0.01) correlation was found between the values of MDA, protein carbonyl-1 and protein carbonyl-2 before the transport.

**Table 2.**
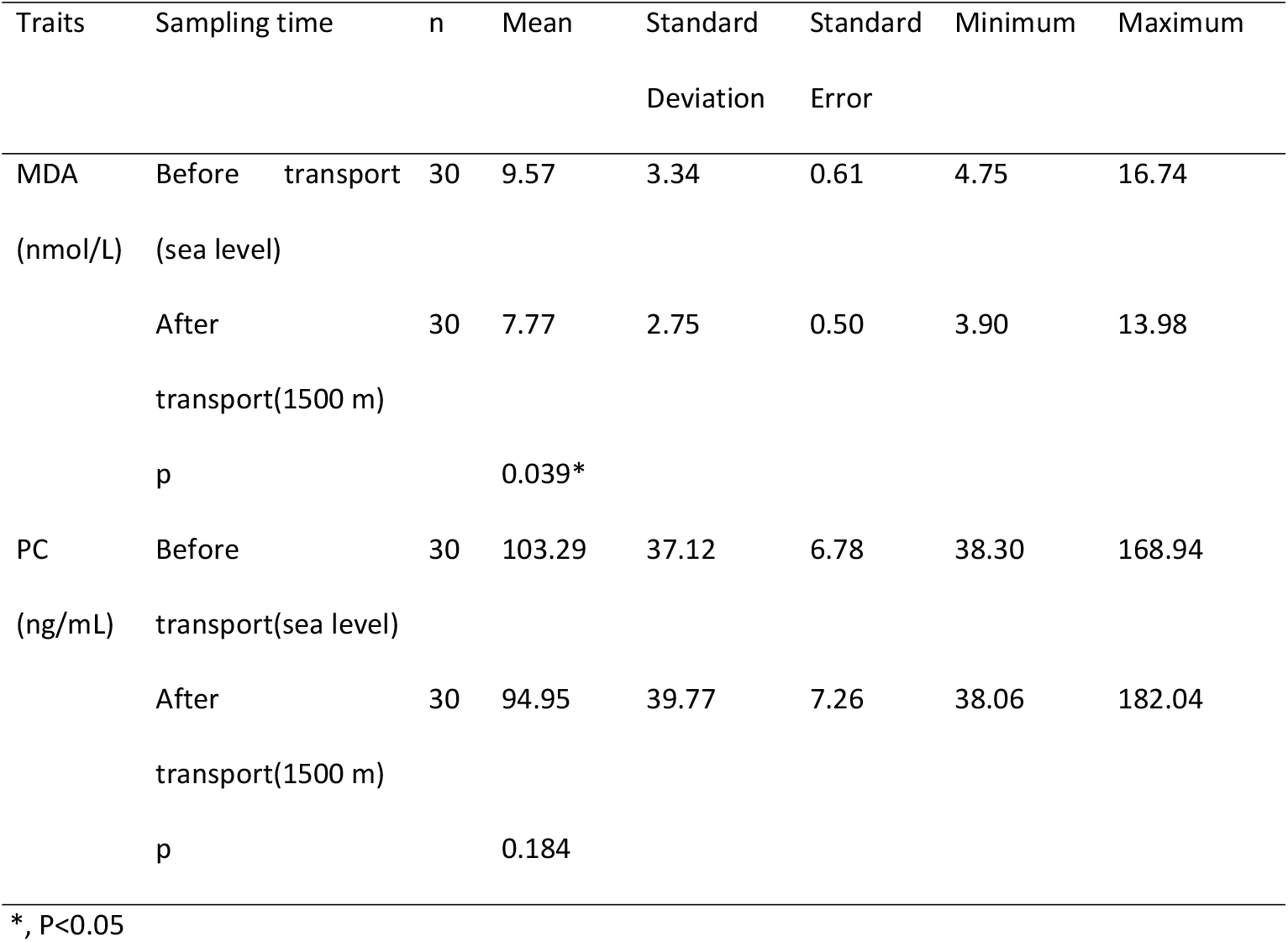
Biochemical parameters at different times in sheep (MDA and Protein Carbonyl)

**Figure 1.**
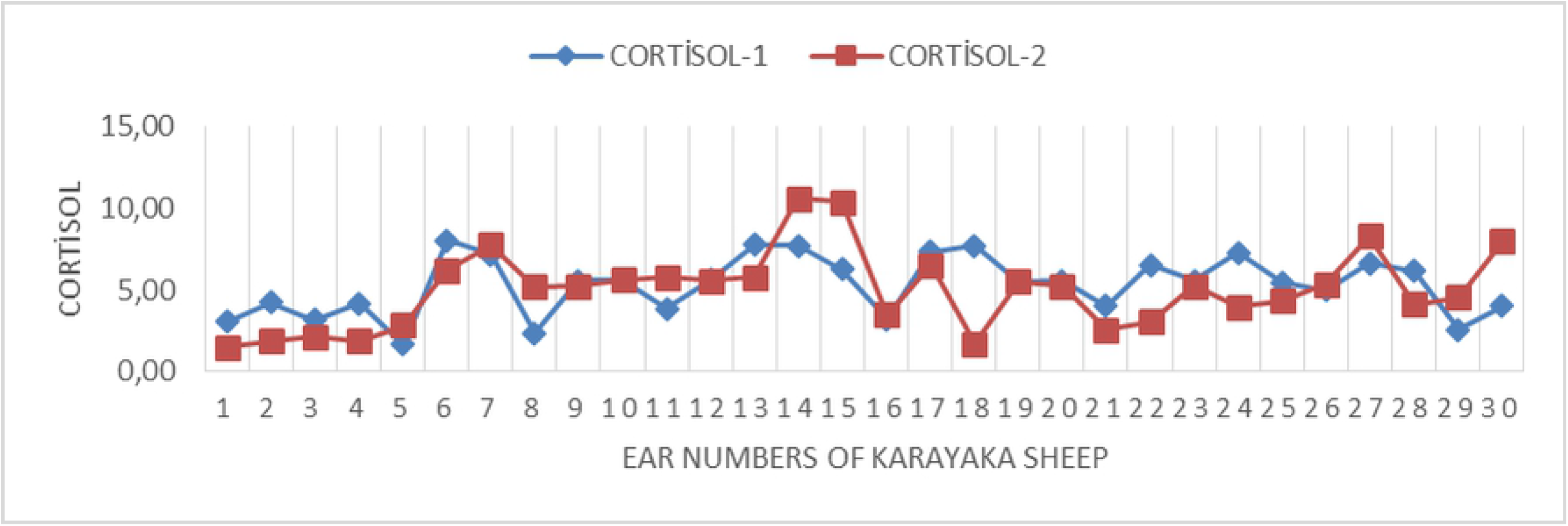
Cortisol hormone change graph in sheep. The changes of cortisol hormone before (sea level) and after transport (1500 m) were graphically given. As seen in Figure 1, in some animals, while cortisol level increases due to transport, it is seen that it is generally close to before transport (sea level). This situation indicates the presence of other stress factors in the pen conditions. Weather heat in the pen may have increased the level of cortisol in sheep.

**Figure 2.**
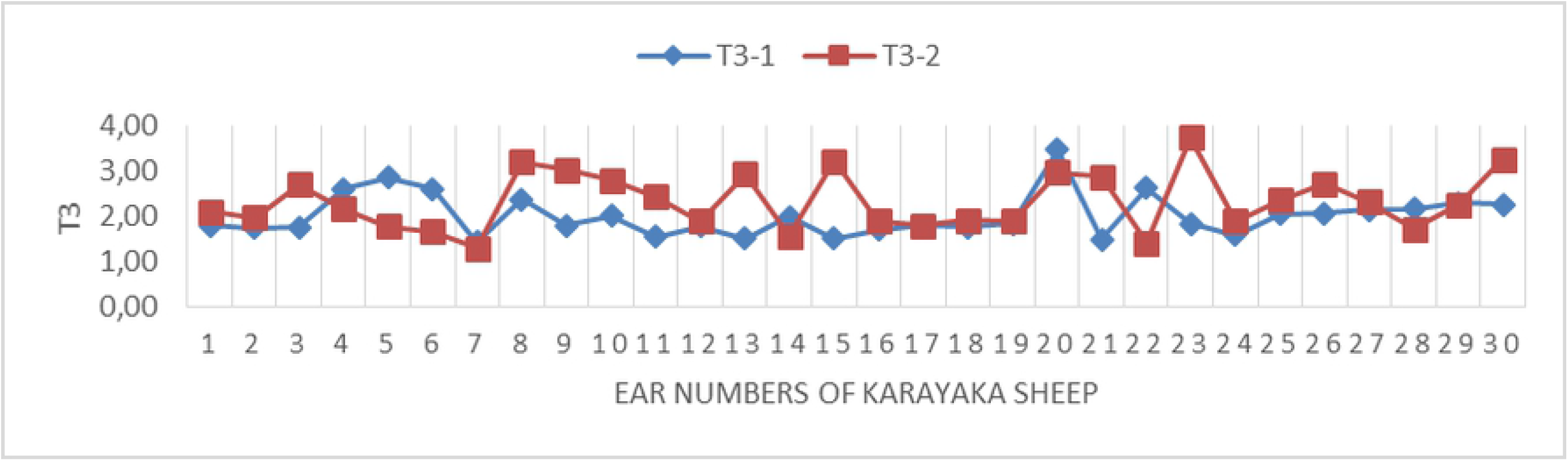
T3 hormone change graph in sheep. The changes of T3 hormone before (sea level) and after transport (1500 m) were graphically given. It is seen that the hormone T3 in Figure 2 increased after transport (1500 m) according to the pen conditions (sea level).

**Figure 3.**
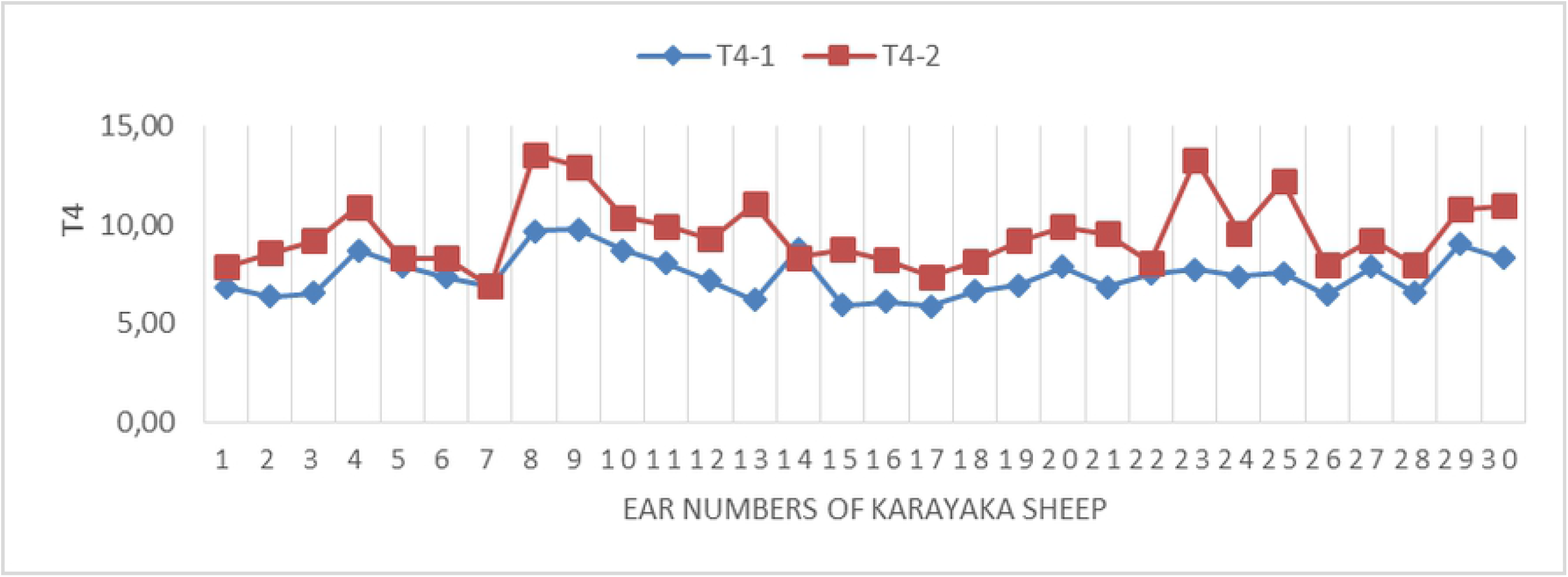
T4 hormone change graph in sheep. The changes of T4 hormone before (sea level) and after transport (1500 m) were graphically given. It is seen that the hormone T4 in Figure 3 increased after transport (1500 m) according to the pen conditions (sea level).

As seen in Table 3, cortisol hormone level was found a significant correlation between the before transport (sea level) and after transport (1500 m). There was a significant (p<0.05) correlation between T3 and T4 hormone before transport and a significant (p<0.01) correlation between T3 and T4 hormone after transport. A very important (p<0.01) correlation was determined between before transport (sea level) T4 hormone and T4 hormone after transport (1500 m).

**Table 3.**
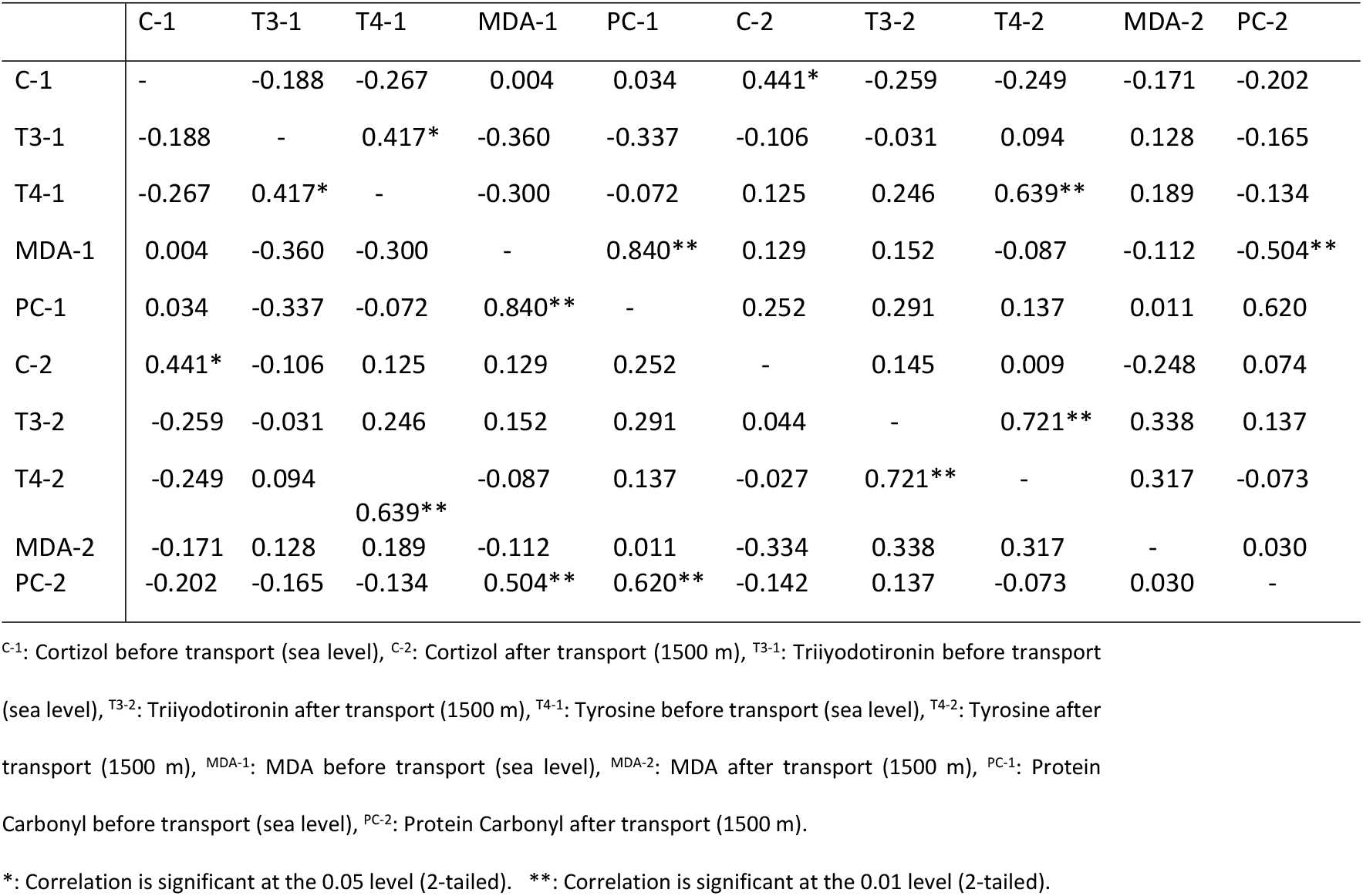
Correlations between hormone and biochemical parameters measured at different times

**Figure 4.**
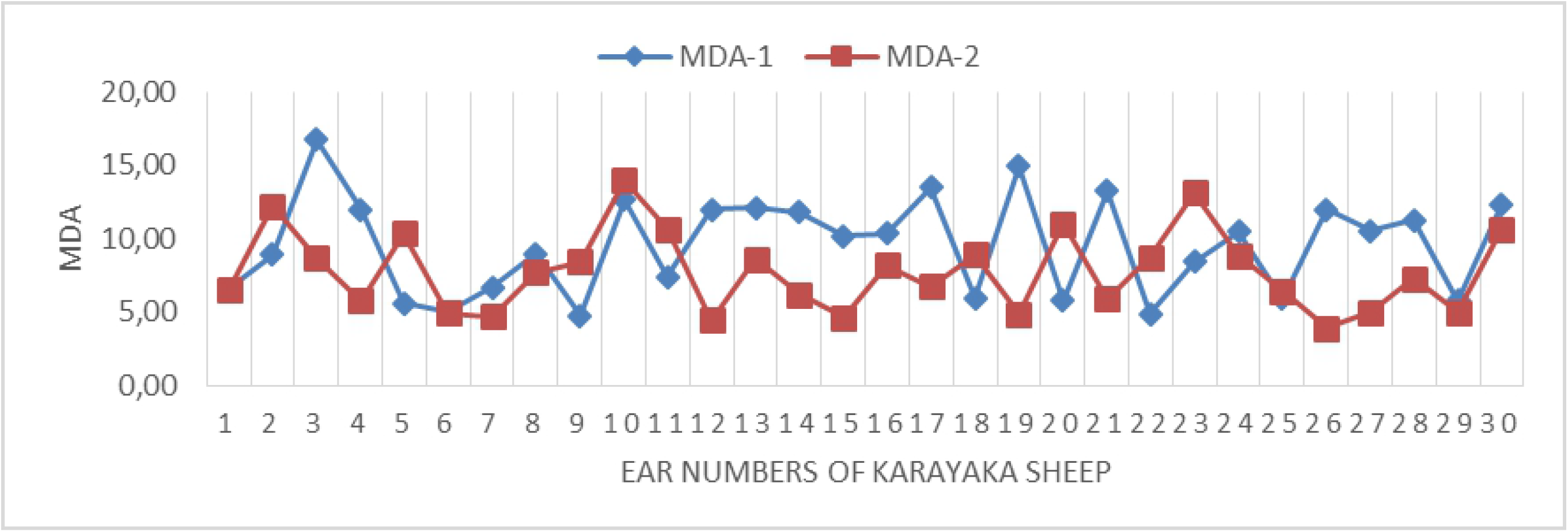
MDA change graph in sheep. The changes of MDA before (sea level) and after transport (1500 m) were graphically given. Figure 4 shows that the MDA value, which is an indicator of lipid peroxidation, has significant differences before transport (sea level) and post-transport (1500 m) values. It is seen that the results obtained before transport (sea level) are higher than the findings obtained after transport (1500 m).

**Figure 5.**
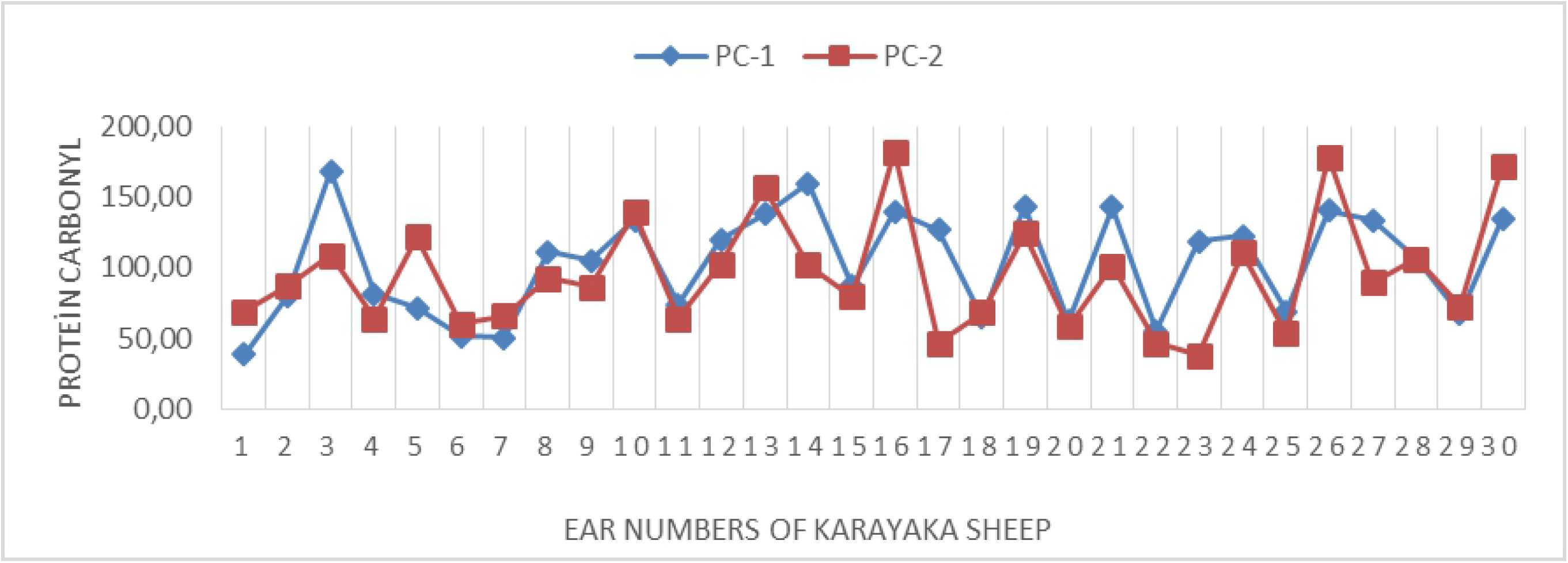
Protein carbonyl change graph in sheep. The changes of protein carbonyl before (sea level) and after transport (1500 m) were graphically given. In Figure 5, it can be seen that the findings obtained before the transport (sea level) are higher than the findings obtained after the transport (1500 m).

## Discussion

### Cortisol hormone

In one study, it was determined that cortisol release increased (P<0.02) in the first 3 hours exposed to transport stress, and then decreased. The cortisol concentration was found to be 5.4 nmol l-1 in the sheep pen [14]. The cortisol (ng/ml) hormone in the before and after transport was determined as 66.421, 255.23 in Morkaraman sheep and 38.954, 104.87 in Imroz sheep, respectively. Cortisol increased after transportation in both races [20]. In Karayaka lambs, cortisol hormone was affected by transport and increased after transport compared to before transport [6]. It has been determined that there is an increase in cortisol levels in the blood as a result of moving the native Morkaraman and Imroz sheep for 75 minutes [21]. In the study, there was no significant change in the level of cortisol with altitude and the transportation of Karayaka sheep. It was determined that is similar the level of cortisol before transport (sea level) with the level of cortisol after transport (1500 m) in Karayaka sheep. It can be said that Karayaka sheep are affected by stress in both cases (Sea level and Perşembe plateo (1500 m)). The insignificant finding of the transportation and altitude effect of cortisol was found different from those reported [6,14,20,21].

### T3 and T4 hormone

T3 (nmol/l) hormone in Iranian fat-tailed sheep (1-2 years old 50-55 kg) in cold environment (4°C), optimum temperature (21°C) and high temperature (40°C), respectively 1.41, 1.26, 0.98; T4 (nmol/l) hormone 59.53, 49.46, 42.44; Cortisol (nmol/l) hormone was determined as 16.56, 10.76, 19.32 [21]. While the hormone T3 and T4 are released too much in the cold environment, it is determined that the release of cortisol in the cold and extremely hot environment increases [21]. It is reported that the general T3 hormone level in Caria sheep is determined as 2.95 ng/ml, it is reported that the air temperature affects the T3 hormone and increases more in the cold [13]. The change of thyroid hormones plays a role in the adaptation to environmental conditions and metabolic balance in living things. T3 and T4 hormone levels decreased as the ambient temperature increased in goats. This is an indicator that metabolism is slowed down and energy production is reduced [22]. In a study conducted in India, it is reported that T3 hormone is low in summer, high in winter, and higher in winter T4 hormone in sheep aged 2-4 [23]. In the study, the findings obtained for the hormone T3 and T4 in Karayaka sheep was compatible with finding [13,20,21,22,23].

### MDA and PC

In the research carried out to determine the effects of transport stress in Akkaraman and Merino sheep, vitamin C was applied prior to transplantation, and it was found that it decreased the level of blood cortisol and did not change the level of lipid peroxidation (MDA) [24]. It is reported that short-term (5 hours) transportation in yearling Akkaraman lambs does not have a significant effect on oxidative stress, and carrying 10 hours or more causes an increase in MDA level [25]. In the study, the findings obtained for MDA in Karayaka sheep was different from the findings [24–25]. It has been determined that cortisol, MDA and PC values are higher in the before transport (sea level) than after transport (1500 m). This situation is thought to have more stress effects before transport (sea level). A high air temperature and humidity may be a factor. In this case, the high temperature in the pen conditions indicates that Karayaka sheep are exposed to heat stress.

We determined the stress effects of transporting and being exposed to altitude difference in a short time (5 hours) in sheep. The results obtained in the study can cause stress in the sheep at the altitude of the destination, even if transportation by vehicle takes a short time. In addition, it was determined that pre-transport conditions of the sheep increased the stress in sheep. With this study, we determined that the barn conditions should be taken into consideration before transportation by vehicle. Stress-reducing measures should be taken in the exposure of sheep to altitude differences and in transportation. Measures that can be taken to reduce the effects of stress are important in terms of increasing the health, productivity and animal welfare of sheep.

In this study, it was determined that sheep raised in closed barns at sea level were exposed to temperature and transport stress and altitude effects. The sheep should be rested during and after the transport. Measures to reduce stress effects should be taken before (sea level) and after transportation (1500 m) (such as adding Ascorbic acid and vitamin E to feed).

## Supporting information

**Figure 1. Cortisol hormone change graph in sheep.**

**Figure 2. T3 hormone change graph in sheep.**

**Figure 3. T4 hormone change graph in sheep.**

**Figure 4. MDA change graph in sheep.**

**Figure 5. Protein carbonyl change graph in sheep.**

## Acknowledgments

This study was supported by the project numbered AR-1808 accepted by the Scientific Research of Ordu University. The authors are grateful to the Scientific Research Projects Commission of Ordu University (AR-1808) for financial support of this study.

## Author contributions

HTC and FAA designed the study. HTC, FAA, MEK and KK performed sampling and animal handling. DUA, TN and SA performed the biochemical analysis. HTC and FAA performed data acquisition and interpretations. HTC and FAA wrote the manuscript. HTC and FAA critically analysed the data and approved the final manuscript. The authors approved the manuscript and agreed to be accountable for liabilities pertaining to the content of this study.

## References

1- Hristov S, Maksimović N, Stanković B, Žujović M, Pantelić V, Stanišić N, Zlatanović Z. The most significant stressors in intensive sheep production. Biotechnology in Animal Husbandry 2012; 28(4):649–658.

2- Cengiz F. Stress generating factors in animals. Journal of the Faculty of Veterinary Medicine 2001; 20:147–153.

3- Hartung J. Effects of transport on health of farm animals. Veterinary Research Communications 2003; 27:525–527.

4- Celi P. The role of oxidative stress in small ruminants’ health and production. Revista Brasileira de Zootecnia 2010: 39:348–363.

5- Piccione G, Casella S, Giannetto C, Bazzano M, Giudice E, Fazio F. Oxidative stress associated with road transportation in ewes. Small Ruminant Research 2013; 112:235–238.

6- Teke B, Ekiz B, Akdag F, Ugurlu M, Ciftci G, Senturk B. Effects of stocking density of lambs on biochemical stress parameters and meat quality related to commercial transportation. Annals Animal Science 2014; 14(3):611–621.

7- Agarwal A, Gupta S and Sharma RK. Role of oxidative stress in female reproduction. Reproductive Biology Endocrinology 2005; 3: 28.

8- McCord JM. Human disease, free radicals, and the oxidant/antioxidant balance. Clinical Biochemistry 1993; 26(5):351–357.

9- Sinatra ST, DeMarco J. Free radicals, oxidative stress, oxidized low density lipoprotein (LDL), and the heart: Antioxidants and other strategies to limit cardiovascular damage. Connecticut Medicine, 1995; 59(10):579–588.

10- Gate L, Paul J, Ba GN, Tew KD, Tapiero H. Oxidative stress induced in pathologies: The role of antioxidants. Biomed Pharmacother 1999; 53(4):169–180.

11- Del Rio D, Stewart AJ, Pellegrini N. A review of recent studies on malondialdehyde as toxic molecule and biological marker of oxidative stress. Nutrition Metabolism Cardiovascular Diseases 2005; 15(4):316–328.

12- Kayali R, Çakatay U. Protein oksidasyonunun ana mekanizmalari. Istanbul University Cerrahpaşa Medical Journal 2004; 35:83–89.

13- Yorulmaz E and Altın T, 2015. The Seasonal Change of Some Physiological Stress Parameters in Sheep. Journal of Adnan Menderes University Agricultural Faculty 2015; 12(2):1–8.

14- Broom DM, Goode JA, Hall SJG, Lloyd DM, Parrott RF. Hormonal and physiological effects of a 15 hour road journey in sheep: comparison with the responses to loading, handling and penning in the absence of transport. British Veterinary Journal 1996; 152(5): 593–604.

15- Altinçekiç ŞÖ and Koyuncu M. Effect of transport conditions on animal welfare. Journal of Animal Production 2010; 51(1):48–56.

16- Fazio E, Ferlazzo A. Evaluation of stress during transport. Veterinary Research Communications, 27 Suppl 2003; 1:519–524.

17- Minka NS, Ayo JO. Physiological responses of food animals to road transportation stress. African Journal of Biotechnology 2010; 9(40):6601–6613.

18- Parraguez VH, Urquieta B, Perez L, Castellaro G, De los Reyes M, Torres-Rovira L, Aguado-Martínez A, Astiz S, González-Bulnes A. Fertility in a high-altitude environment is compromised by luteal dysfunction: the relative roles of hypoxia and oxidative stress. Reproductive Biology and Endocrinology 2013; 11(24):1–12.

19- Adili N, Melizi M. The effect of age, sex and altitude on the morphometry of red blood cells in small ruminants. Journal of Animal Science Advances 2013; 3(1):27–32.

20- Nazifi S, Saeb M, Rowghani E, Kaveh K. The influences of thermal stress on serum bio-chemical parameters of Iranian fat-tailed sheep and their correlation with triiodothyronine (T3), thyroxine (T4) and cortizol concentrations. Comparative Clinical Pathology 2003; 12:135–139.

21- Ekiz B, Ergül Ekiz E, Yalçintan H, Yilmaz A, Koçak Ö, Güneş H. Effect of ram-ewe mixed transportation on certain welfare parameters in red Karaman and Imroz sheep. Journal of Faculty of Veterinary Medicine, Istanbul University 2013; 39(2):155–167.

22- Koluman Darcan N, Daşkıran İ, Şener B. The heat strees effect on T4 (thyroxin), T3 (triiodothyronine), cortisol hormones of goats in rearing extensive systems. Journal of Tekirdag Agricultural Faculty 2013; 10(3):29–36.

23- Sawankumar DR, Vasava AA, Pathan MM, Madhira SP, Patel YG, Pande AM. Effect of season on physiological, biochemical, hormonal, and oxidative stress parameters of indigenous sheep. Veterinary World 2017; 10(6):650–654.

24- Avci G, Küçükkurt İ, Fidan F, Eryavuz A, Aslan R, Yılmaz D. The Influence of vitamin C and xylazine on cortisol, lipid peroxidation and some biochemical parameters in transported sheep. Firat University Veterinary Journal of Health Sciences 2008; 22(3):147–152.

25- Çetin E, Çetin N, Küçük O. The effect of road transport on oxidant-antioxidant system in yearling lambs. Atatürk University Journal of Veterinary Sciences 2011; 6(2):103–109.

